# The network makeup artist (NORMA-2.0): Distinguishing annotated groups in a network using innovative layout strategies

**DOI:** 10.1101/2022.03.02.482621

**Authors:** Evangelos Karatzas, Mikaela Koutrouli, Fotis Baltoumas, Katerina Papanikolopoulou, Costas Bouyioukos, Georgios A. Pavlopoulos

## Abstract

**Motivation:** Network biology is a dominant player in today’s multi-omics era. Therefore, the need for visualization tools which can efficiently cope with intra-network heterogeneity emerges.

**Results:** NORMA-2.0 is a web application which uses efficient layouts to group together areas of interest in a network. In this version, NORMA-2.0 utilizes three different strategies to make such groupings as distinct as possible while it preserves all of the properties from its first version where one can handle multiple networks and annotation files simultaneously.

**Availability:** The web resource is available at http://norma.pavlopouloslab.info/

The source code is freely available at https://github.com/PavlopoulosLab/NORMA

## 1 Introduction

Network visualization is important to capture patterns and understand the associations among various biomedical entities coming from different repositories (Koutrouli *et al*., 2020; Baltoumas *et al*., 2021). To this end, several visualization tools have been proposed (Gehlenborg *et al*., 2010). However, despite their richness in the interactivity and variety of analysis options they offer (e.g., Cytoscape’s apps (Saito *et al*., 2012)), only few of them such as Cytoscape (Shannon *et al*., 2003) or Arena3D^web^ (Karatzas *et al*., 2021) can cope with node heterogeneity. What is more, most of these tools focus mainly on interactivity, layout, and network exploration and lack efficiency in effectively visualizing network annotations. To alleviate this issue, we developed the Network Makeup Artist (NORMA) application. NORMA is able to handle multiple networks and annotations simultaneously and produce high-quality, publication-ready visualizations. It also supports interactive network annotation visualization, topological analysis and automated community detection with various algorithms.

In this article, we present NORMA-2.0, a major update of the original NORMA application (Koutrouli *et al*., 2021) which now incorporates three different layout strategies to make annotated groups or areas of interest in a network more visually distinct. This way, NORMA-2.0 offers options for more appealing representations and enables a more targeted knowledge extraction and storytelling.

## 2 Methods

### 2.1. General functionality

NORMA is a handy web tool for interactive network annotation, visualization, and topological analysis through which users are able to handle multiple networks and annotations simultaneously. It offers group highlighting with the use of shaded areas (convex hulls) or pie-chart-like nodes while it comes with a functionality for directly comparing several networks topologically as well as different group annotations for the same network. In addition, users can perform clustering analysis on-the-fly and highlight nodes using any custom color scheme (e.g., expression values).

### 2.2. Implementation

In this version, NORMA-2.0 utilizes three different strategies to visually separate annotated areas of interest in a network in combination with established layout algorithms, offered by *igraph* (Gabor Csardi and Tamas Nepusz, 2006). NORMA-2.0 is written in R/Shiny. Network visualization is offered by d3.js. The 3D network and convex hull visualizations are offered by plotly.

***Strategy 1 – Virtual nodes:*** As in the original version, NORMA-2.0 introduces one virtual node per group which behaves as a hub. Upon creation, edges with heavy weights are assigned to this node, linking it with all nodes from the same group. Then, any traditional layout algorithm can be utilized until it converges. The difference compared to directly applying a layout algorithm on the network is that the virtual nodes will attract the group-specific nodes as if they were parts of the network. After completing the layout execution, all virtual nodes are removed (Figure 1A).

**Figure 1.**
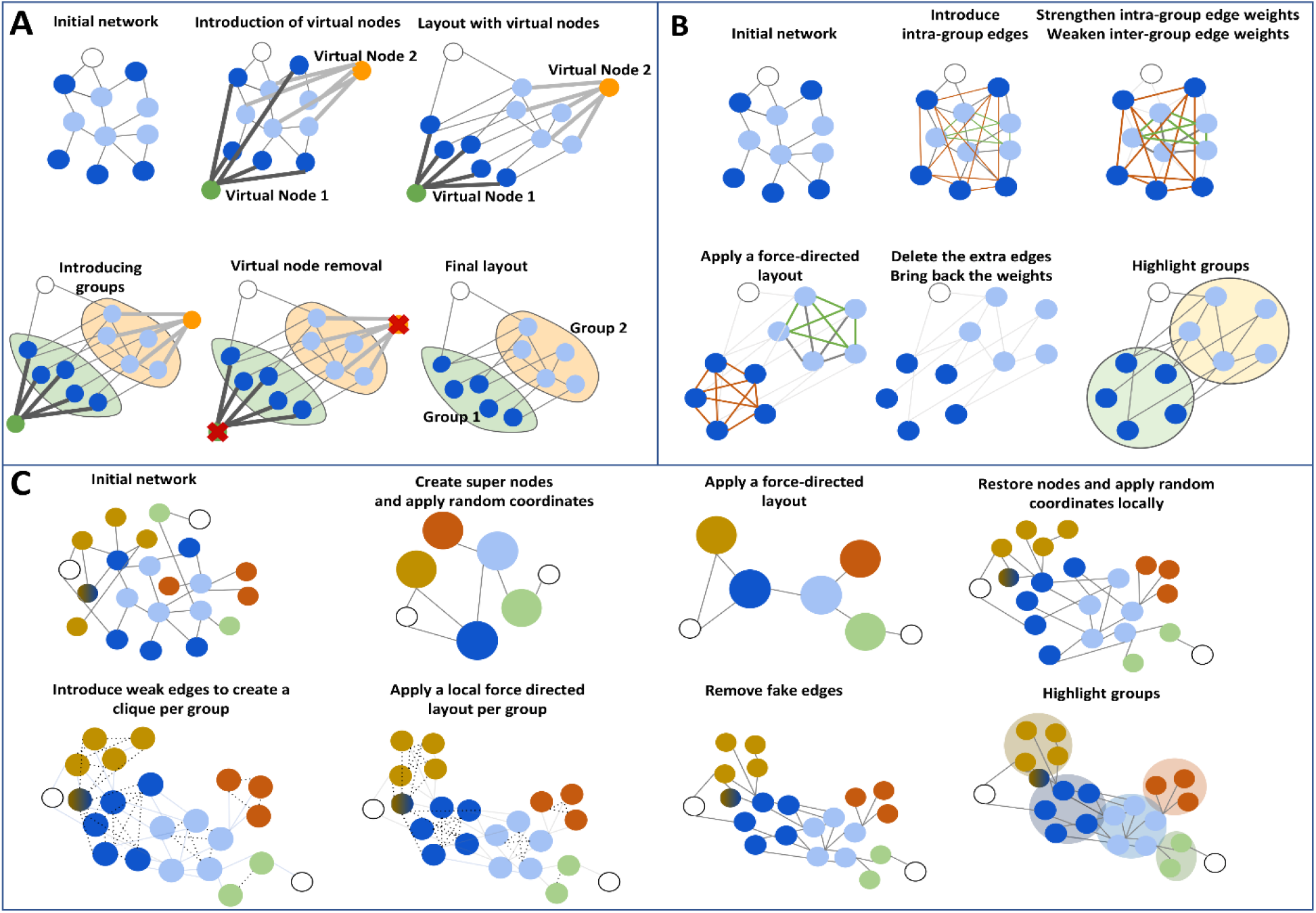
Three different layout strategies to highlight areas of interest within a network and make them as distinct as possible.

***Strategy 2 - Gravity:*** Here, NORMA-2.0 introduces intra-group edges where necessary to generate clique-like subnetworks (all-vs-all connections). As a second step, the intra-group edge weights are significantly increased whereas the inter-group edge weights are simultaneously decreased. Then, any of the offered layouts can be applied to adjust node coordinates. The introduced edges and weights only exist for the calculation of the layout coordinates, and do not carry over to the final visualized network (Figure 1B).

***Strategy 3 – Super nodes:*** In this scenario, NORMA-2.0 introduces ‘super-nodes’ to represent each of the uploaded annotation groups. Then, it connects these groups with edges which correspond to the connections from the initial network. For example, if node A belonging to the annotation group 1 is connected to node B from group 2, then the group 1 super-node will also be connected to the group 2 super-node. Notably, the network becomes significantly smaller both in terms of node and connection numbers. In a second step, any of the available layouts can be applied on the ‘super-network’. Upon layout convergence, all centroid coordinates of these super-nodes as well as the coordinates of no-group nodes can be further repelled according to a user-defined input. Then, all initial nodes will be placed around their respective super-nodes according to a second user-selected local layout choice. If a node belongs to more than one group, the average values for their (x, y) coordinates are used for the final visualization (Figure 1C).

### 2.2 Application to the *Drosophila* Tau network

To showcase the capabilities of NORMA-2.0, we use the *Drosophila* Tau protein-protein interaction network (Figure 2), based on experimental evidence from (Papanikolopoulou *et al*., 2019) and described in detail in (Koutrouli *et al*., 2021). Briefly, the network consists of 358 differentially expressed proteins from *Drosophila* Null-Tau mutants connected by 3176 edges, with each node colored based on its expression evidence (green for up-regulated, red for down-regulated). Network clustering was performed using the Louvain algorithm (Blondel *et al*., 2008) based on its topology, and the resulting communities are modeled as network groups. Compared to the original network or the virtual nodes strategy (Figure 2A-B), the application of NORMA-2.0’s novel layout modification strategies (Figure 2C-D) can result in more comprehensive views, in which the different community groups are easier to detect and visualize. Especially in the case of using supernodes (strategy 3), the ability to use a local layout alongside the global network layout produces a well-defined, aesthetically pleasing visualization.

**Figure 2.**
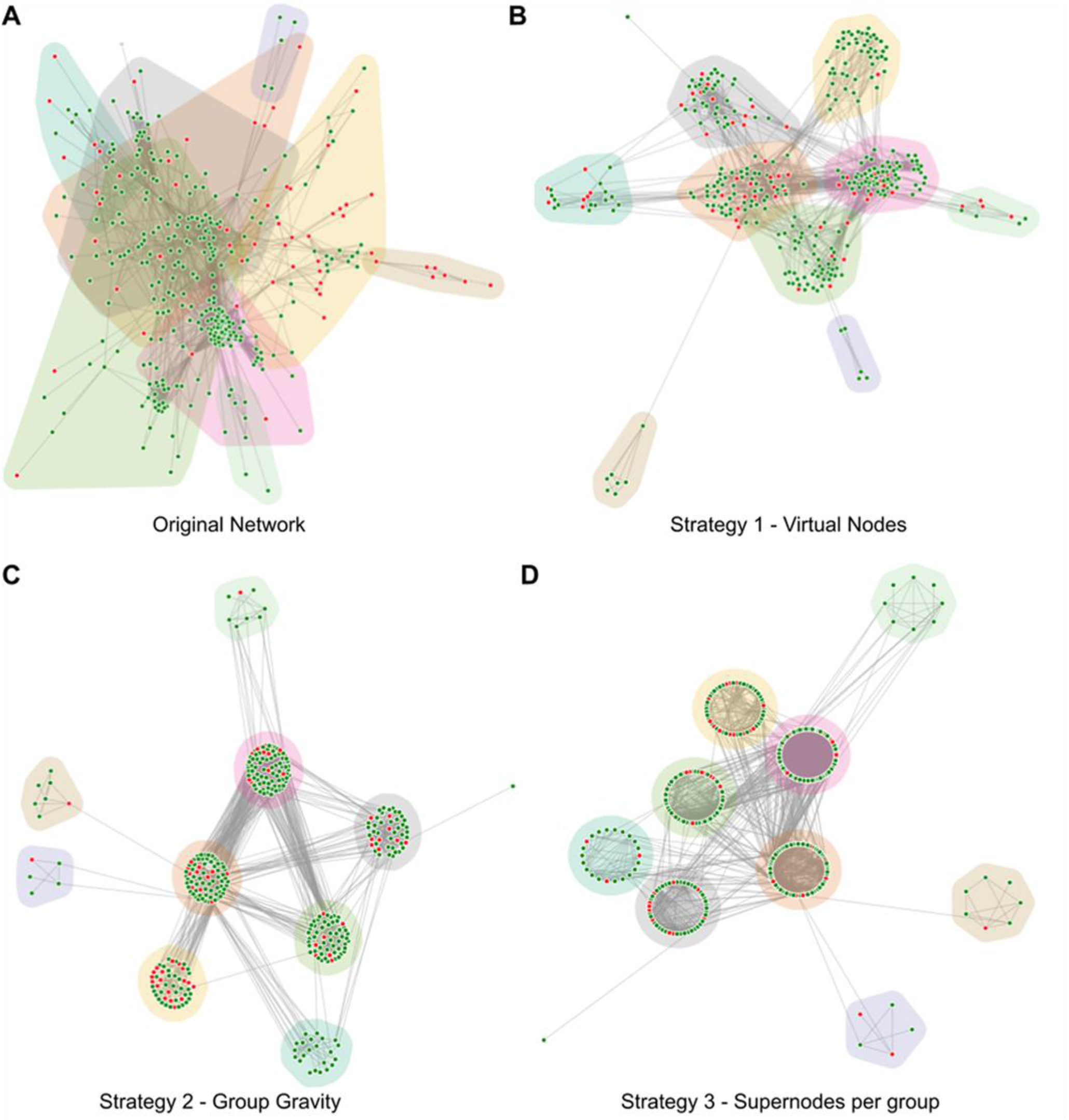
Visualization of the *Drosophila* Tau network in NORMA-2.0, using a force-directed layout. A) The original network with no layout modifications, B), Application of strategy 1 (virtual nodes), C) Application of strategy 2 (group gravity), D) Application of strategy 3 (supernodes). A force-directed layout is used for the global network, while a local circular layout is used for each group.

## 3 Discussion

We present NORMA-2.0, an update to the original NORMA network visualization platform that implements novel visualization strategies and improvements to its performance and utility. The main strength of NORMA-2.0, compared to its previous iteration and other network visualization platforms is ease of use, as well as the automated generation of appealing network visualizations based on user-provided annotations. While complex network views, such as those described, can be rendered with various tools (e.g., through Cytoscape’s advanced Filtering tab), their creation can be time consuming and complex, particularly for inexperienced users. Conversely, NORMA-2.0 can produce high-quality, publication-ready network views with minimal input in the form of simple, easy to create annotation files. Overall, we believe that NORMA-2.0 is a handy tool for fast and high-quality, complex network visualization, designed for beginners as well as experts.

## ACKNOWLEDGEMENTS

We would like to acknowledge support from the Hellenic Foundation for Research and Innovation (H.F.R.I) under the “First Call for H.F.R.I Research Projects to support faculty members and researchers and the procurement of high-cost research equipment grant”, Grant ID: 1855-BOLOGNA. GAP was also supported by the project ‘The Greek Research Infrastructure for Personalised Medicine (pMedGR)’ (MIS 5002802), which is implemented under the Action ‘Reinforcement of the Research and Innovation Infrastructure’, funded by the Operational Program ‘Competitiveness, Entrepreneurship and Innovation’ (NSRF 2014-2020) and co-financed by Greece and the European Union (European Regional Development Fund). Finally, this article was partially funded by the Greek matching funds for the Marie Sklodowska-Curie Individual Fellowships Grant—MSCA-IF-EF-CAR (Grant ID: 838018-H2020-MSCA-IF-2018).

